# DREAMER-S: Deep leaRning-Enabled Attention-based Multiple-instance approaches with Explainable Representations for Spatial biology

**DOI:** 10.1101/2025.10.02.679953

**Authors:** M. Rifqi Rafsanjani, Alison Dooney, Rahul Suresh, Alice C. O’Farrell, Monika A. Jarzabek, Liam Shiels, Annette T. Byrne, Jochen H. M. Prehn, Aidan D. Meade

## Abstract

Identifying image features that associate strongly with diagnostic or prognostic classes in large-scale, multi-channel spatial imaging is challenging without pixel-level annotations. We present DREAMER-S, an attention-based multiple-instance learning (MIL) framework that, using only image- or slide-level labels, learns spatial features within 3D imaging hypercubes that are most informative for downstream classification. We demonstrate DREAMER-S on Quantum Cascade Laser infrared (QCL-IR) tissue imaging, where attention weights are rendered spatially to highlight class-relevant spectral instances without manual annotation. Because the MIL attention layer assigns interpretable importances to spatial instances, the method is broadly transferable to spatial-biology applications that require instance-level filtering to focus towards salient regions of interest in high-content datasets.

We further evaluate DREAMER-S on a chemotherapy-response task in a colorectal cancer patient-derived xenograft (PDX) model. After tuning, DREAMER-S separated spectral instances from a chemo-sensitive PDX (CRC0344) and a less responsive PDX (CRC0076) with an F1 score of ∼0.95.

To validate explainability, we linked model saliency to cellular physiology, observing that, (i) unsupervised UMAP embeddings of high-attention spectra stratified samples by treatment (chemotherapy, apoptosis sensitizer, combination, vehicle), and (ii) selected spectral markers correlated with pro-apoptotic proteins measured independently in the same PDX system. Together, these results support a mechanistic link between spectral signals and apoptosis pathways and position DREAMER-S as an efficient, interpretable approach for analysing high-content spatial-biology imaging datasets.

## 1. Introduction

Multiple Instance Learning (MIL) is an innovative machine learning approach designed to address challenges associated with weakly labelled data, where class labels are assigned to collections of instances, known as bags, rather than to individual instances themselves. A bag is labelled positive if at least one of its instances is positive, while it is labelled negative only if all instances are negative (1). This approach is particularly beneficial in scenarios where labelling is expensive or impractical, allowing for the classification of bags based on the presence of positive instances within them (2,3). Hence, it is a common strategy in medical imaging particularly in digital pathology, to analyse whole slide images (WSIs) which are subdivided into smaller patches so that a deep neural network model can be trained for specific purposes such as detecting epithelial cells, grading tumours, and identifying other histopathological features (reviewed systemically in (4)). MIL can therefore facilitate the simultaneous feature localisation and classification of disease in a computationally-efficient manner in large images, addressing the inherent challenges posed by the complexity and size of WSIs (5).

Mid-infrared (IR) spectroscopy allows for the identification of specific molecular signatures associated with various pathological conditions using comprehensive biochemical profiles of tissues through the analysis of the absorption of infrared light. This is because biomolecules absorb in the mid-IR (4000 cm^−1^ to 400 cm^-1^) producing an absorbance spectrum which can be treated as a unique biochemical fingerprint. This fingerprint can be used to differentiate between normal and tumour tissues (6), identify spectral signatures indicative of endometriosis (7), elucidate the chemical composition of lipid droplets by size in liver tissue and endothelial cells (8), amongst other applications. IR spectroscopy is therefore an attractive approach for histopathological analysis due to its non-invasive, label-free, objective, and quantitative nature enabling clinically informed decisions without the need for extensive sample preparation or labelling (9).

In a previous study, from which the present study stems, (10), the response of two patient derived xenograft (PDX) mouse colon cancer models were studied (CRC0344 and CRC0076), which differed in their responsiveness to chemotherapy. The CRC0344 model was found to be sensitive to 5-fluorouracil (5-FU)-based chemotherapy, while the CRC0076 model was less responsive to chemotherapy but responded with tumour regression by addition of the apoptosis sensitiser and BCL-2 antagonist, ABT-199. Traditional imaging methods such as 18F-Fluorodeoxyglucose positron emission tomography/computed tomography (18F-FDG-PET/CT) were demonstrated to have potential as early response biomarkers (10). The central hypothesis of the present study was that applying a deep learning MIL (DL-MIL) approach to the rich biochemical data from IR chemical imaging of the PDX colon cancer tissue would uncover unique spectral biomarkers capable of discriminating each PDX-model on their sensitivity to chemotherapeutic treatment.

Modern IR imaging microscope systems employing focal plane array (FPA) detectors, or similar, allow the generation of full spectral information spatially across a sample, such that the output “chemical” images are termed hyperspectral data cubes. The challenge with using these images within a deep learning modelling approach in the absence of dimensionality reduction is the sheer size of each image in data terms. For example, a single chemical image generated by a 256 × 256 FPA array capturing 217 wavenumbers per spectrum will contain 14,221,312 datapoints. Comparatively a typical image size within the CIFAR10 dataset commonly used as a training dataset for deep learning only contains 32 x 32 spatial pixels, each with 3 RGB channels, or 3072 datapoints (11). Therefore, our hypothesis is that the use of a deep learning-MIL framework could represent a practical approach in this context, in terms of both computational efficiency and identification of spectral region of interests (ROIs) with biochemistry that are linked to the classification target (in this case identification of chemotherapeutic resistance or sensitivity).

A number of recent studies have employed weakly supervised approaches with vibrational spectroscopic data for the analysis of both imaging or point-spectral data. For example, MIL and Support Vector Machine (SVM) algorithms have been applied to Raman spectroscopy for the detection of COVID-19 in saliva samples (12), while MIL and Random Forest (RF) algorithms have been utilised in IR studies to identify aberrant tissue in mouse livers (13). Research by Shi et al (14) and Phan et al (15) has demonstrated the potential of the MIL approach with CNNs for the identification of regions of interest with spectral images containing signatures of drug fingerprints and microplastics, respectively. However, to our knowledge no previous study has employed an integrated MIL and deep learning (MIL-DL) pipeline with chemical imaging data and explainability approaches to identify spectral biomarkers driving systemic responses to treatment within heterogenous tissues.

In this paper, a MIL-DL framework was employed to train and optimise a deep learning model to classify chemotherapeutic sensitivity from quantum cascade laser (QCL) IR chemical imaging data, and candidate spectral biomarkers driving the classification were extracted using explainability approaches. Model architecture and model hyperparameter settings (number of layers, the depth of the network, the number of hidden nodes, the residual connection within the network and the optimiser’s learning rate) were tuned using two strep grid-parameter search strategy. Then candidate spectral biomarkers identified by the model were extracted using the attention mechanism, and SHAP values within the best performing model. The correlation between treatment group and top attended spectral biomarkers were also further explored using Uniform Manifold Approximation and Projection (UMAP) clustering.

While the DREAMER-S framework is demonstrated in the domain of spectral histopathology, it acts as a data-reduction approach and ROI-filter that can be included within high-content spatial biology pipelines for diverse applications, e.g., Raman/CARS, mass spectrometry imaging, multiplexed immunofluorescence, and potentially spatial transcriptomics or proteomics, where manual pixel-level annotation is infeasible or unavailable.

## 2. Experimental Section

### PDX-derived colon cancer tissue sections

We utilised PDX samples generated in a prior study (10). The original PDX tissue material was derived from metastatic CRC (mCRC) patient liver lesions, and established by Bertotti et al. (16). All patients provided informed consent. Samples were procured under the approval of the Review Boards of the Institutions (PROFILING protocol No. 001-IRCC-00IIS-10). All animal procedures were approved by the Health Products Regulatory Authority (HPRA) (#AE18982-P099) and the University College Dublin Animal Research Ethics Committee (AREC) (#AREC-16-11). Two PDX models were selected based on their predicted sensitivity to chemotherapy, one sensitive to 5-FU-based chemotherapy (labelled as CRC0344) and another sensitised by ABT-199 (labelled as CRC0076). Animals bearing either subcutaneous CRC0076 or CRC0344 tumours received one of the following treatment regimens: (i) FOLFOX (all constituent drugs delivered IP, once weekly (on day 3 of each cycle), 5-FU (40 mg/kg in PBS) + Folinic Acid (13.4 mg/kg in PBS) followed two hours later by OX (2.4 mg/kg in 5% glucose/water (v/v)), (ii) ABT-199 (oral gavage, once daily at 100 mg/kg, dissolved in 60% phosal, 50 propylene glycol (PG), 30% polyethylene glycol (PEG) 400 and 10% ethanol), (iii) FOLFOX with ABT-199 (dosed as described above) and (iv) a vehicle control (all diluent vehicle solutions), resulting in eight treatment classes. Researchers undertaking the animal studies were blinded to the expected sensitivity of the PDX models. At the end of the study (i.e. after 4 weeks of treatment or when the study reached humane end point [based on a scoring system accounting for tumour size and animal wellbeing]), tumours were excised, rinsed twice in Dulbecco’s-(D)PBS and fixed in 4% formaldehyde for 48 h and embedded in paraffin.

### Acquisition of chemical imaging data

A 5 µm-thick tissue section was prepared and mounted on a calcium fluoride (CaF_2_) slide for analysis. To retain all tissue the section was not subjected to any chemical dewaxation. Spectroscopic measurements were carried out using a Daylight Spero-QT 340 Quantum Cascade Laser (QCL) infrared microscope operating in transmission mode. Hyperspectral chemical images (HCIs) were captured at a low magnification (0.3 NA) across the wavenumber range 952– 1800 cm^−1^. The generated HCIs comprised a spatial area of 480×480 pixels and a spectral depth of 213 wavenumbers.

### Spectral pre-processing

Following data acquisition, a multi-step pre-processing pipeline was applied to the raw spectral data using custom scripts written in Python (version 3.10). The primary objective of these steps was to correct for artefacts and to isolate the spectra originating specifically from the tissue. First, a rubber-band baseline correction was applied to each spectrum using the Pybaselines library (version 1.2) (17). Then, a thresholding value of 0.1 was applied at the absorbance intensity of the Amide I peak (located at approximately 1654 cm^-1^) to distinguish tissue regions from the background where any spectrum below the thresholding value was classified as non-tissue and subsequently replaced with zero-vectors. Finally, vector normalisation was performed on the remaining tissue spectra using Scikit-Learn module (version 1.7.0) (18). The full dataset for this study comprised 40 hyperspectral images. Each image, containing 213 wavenumbers and 230,400 individual spectra, represents over 49 million data points in each image. The complete pre-processing workflow, from raw data to corrected spectra, is visually summarised in **Figure 1**a.

**Figure 1.**
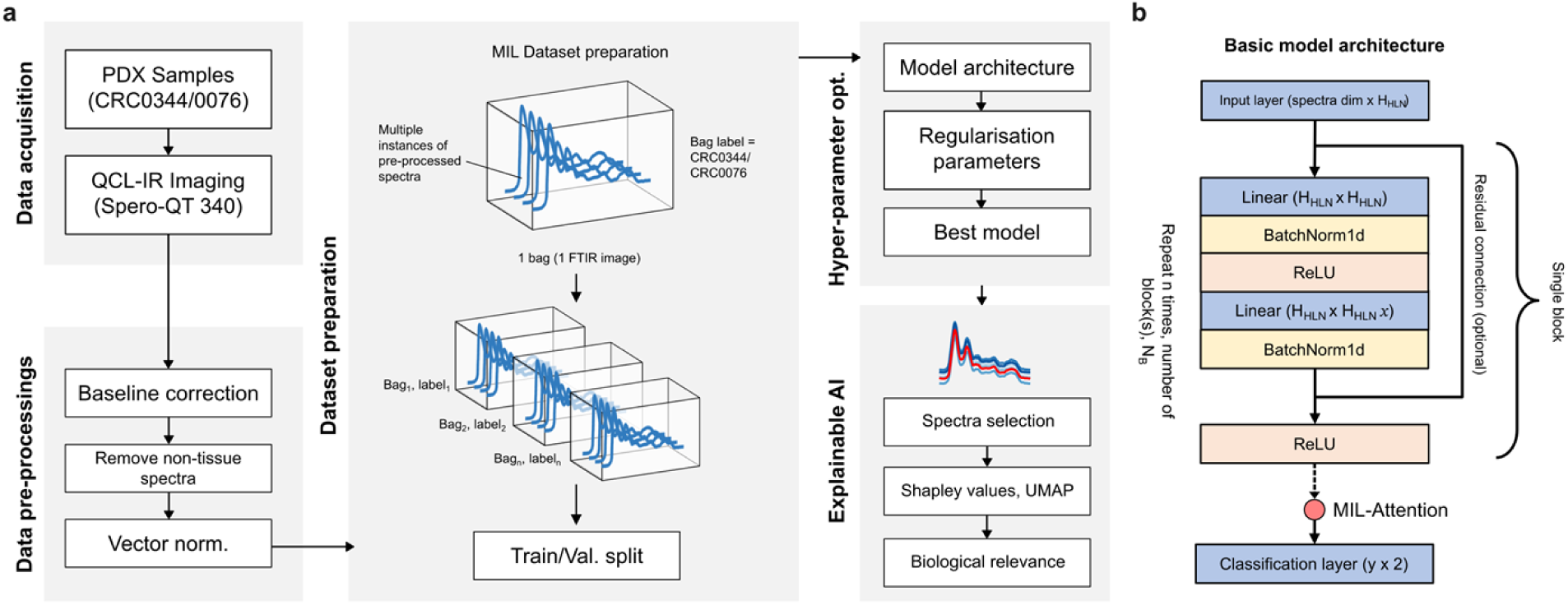
Overview of the workflow of the multiple-instance learning (MIL) approach applied to hyperspectral chemical image data. (a) A summary of the preprocessing steps for chemical images, including preparation procedures prior to training the deep learning model and optimising hyperparameters. (b) The skeletal structure of the linear model architecture utilised in this study, which is loosely inspired by the ResNet framework.

### Multiple instance learning dataset preparation

For each chemotherapy sensitivity group (CRC0076 and CRC0344), the hyperspectral image dataset was manually partitioned into equal subsets (50%:50%) for training and validation. Each subset comprised of 20 hyperspectral images: 10 hyperspectral images for each of CRC0076 and CRC0344 (Supplementary section **Table S 1**). Both training and validation datasets contained a representative distribution of the treatment conditions described earlier (FOLFOX, ABT-199, FOLFOX with ABT-199 and a vehicle control) and each group was separated by the identity of individual mice to prevent data leakage during model training and testing. In our multiple instance learning (MIL) approach, each hyperspectral image was treated as a bag consisting of multiple spectral instances (230,400 individual spectra in each bag). Each bag was assigned a label corresponding to its respective chemotherapy sensitivity group (label 0 for CRC0076 and label 1 for CRC0344) for the classification model training task. During the MIL training, the whole bag (chemical image) was fed through the network for a single prediction class.

### Deep learning model architecture

Deep learning model architectures were developed using Pytorch (version 2.4.0) (19) with their design loosely inspired by the Residual-Net (ResNet) block where each block consists of repeating linear, batch normalisation and ReLU activation layers (see **Figure 1**b) (20). An optional residual connection was introduced before the first linear layer that connected the final ReLU activation layer in each block. An input layer was also introduced prior to the first block to allow the input of spectral data and a final linear layer was introduced for multi-class classification. Following the sequence of blocks, and preceding the final classification layer, a MIL-attention mechanism was applied based on the implementation from (21). The purpose of this mechanism is to produce a single, representative embedding for the entire input “bag” by weighting the significance of each instance. This allows the model to identify which instances are most influential for the final prediction. Subsequently, a final linear layer uses this attention-pooled embedding for multi-class classification. The MIL attention mechanism can be summarized as follows:

- let *H={h_1_, h_2_, …, h_K_}* represent a bag containing *K* instance embeddings, where each instance *h_k_* ∈R^M^ is an *M*-dimensional real-valued feature vector representing an individual instance.

- compute a weighted average, *z*, of these instances to form a single bag-level representation:

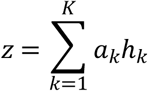

The attention weights, *a_k_*, are calculated using a small neural network and normalized across all instances in the bag via the softmax function. This ensures that the weights sum to 1 and allows the model to learn the relative importance of each instance:

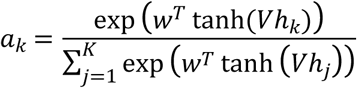

where *V∈R^L×M^* and *w∈ ^RL^* are the learnable weight parameters of the attention network. This formulation enables the model to focus selectively on the most informative instances within each bag, thereby enhancing the overall discriminative power of the bag-level representation in weakly supervised learning settings.

### Hyperparameter optimisations

To determine the optimal model configuration for our hyperspectral data, we conducted a systematic grid search of key hyperparameters. This optimisation process focused on two main areas: 1) the model’s fundamental architecture and 2) its learning rate parameter. The main goal was to maximise classification performance while balancing model interpretability, maintaining computational efficiency, and mitigating the risk of overfitting. The architectural search explored several core parameters. First, network depth was varied by adjusting the number of sequential blocks (N_B_), evaluating shallow (N_B_=1) and deeper (N_B_=2 or 3) networks. Second, the network width was modified by changing the number of nodes (N_HLN_) within the fully connected layers, with values ranging from 64 to 256. We also investigate the network’s expansion strategy by assessing two distinct approaches. In the ’expanding’ configuration, the number of nodes was doubled in each successive block, a design intended to capture increasingly complex and hierarchical features. Conversely, the ’shrinking’ configuration halved the number of nodes per block, progressively reducing the network’s complexity. Furthermore, the inclusion of residual connections was tested as an additional parameter to assess their potential to improve gradient flow and overall model performance, particularly in deeper architectures.

The optimisation was performed in a two-stage process. Initially, the optimal architectural parameters were identified using a default learning rate (1x10^-3^). Once the best-performing architecture was established, its specific learning rate for the Adam optimiser was then fine-tuned. Throughout this process, Cross-Entropy was employed as the loss function, while the F1-score served as the primary metric for evaluating model performance during the training and validation phases. To ensure the robustness of our findings, each hyperparameter combination was trained for 20 epochs and the process was repeated at least three times with different random seeds. The final selection was based on the parameter set that yielded the highest average F1-score on the validation dataset. A comprehensive summary of the hyperparameters and their tested values is provided in **Table 1**. Each hyperparameter set was evaluated across at least three independent runs, and the configuration with the highest F1-score on the validation set was selected. This chosen configuration was then retrained for 50 epochs, and the best-performing model checkpoint was saved for further analysis.

**Table 1.** List of hyperparameters and their values for the grid search hyper parameter model optimisation.

| Hyperparameters | Descriptions | Values |
| --- | --- | --- |
| Layer's depth (number of blocks, $N_B$) | Depth of the neural network | 1, 2, 3 |
| Number of nodes in first hidden layer, $N_{HLN}$ | Number of nodes of the first linear layer | 64, 128, 256 |
| Model's expansion type | Either to expand (i.e: from 64 to 128 for double) or reduce (from 64 to 32 for half) | double, half |
| Residual connections | Option to add residual connection for deeper networks | True, False |
| Optimiser's learning rate | The rate at which a model learns by adjusting the step size during training | $1 \times 10^{-1}$, $1 \times 10^{-3}$, $1 \times 10^{-5}$ |

**Table 2.** Training results for learning rate parameter for both validation and training dataset over three trial run with standard deviation based using the optimum model architecture parameter.

| Learning rate | Loss values |  | F1 Scores |  | Accuracy |  |
| --- | --- | --- | --- | --- | --- | --- |
|  | Val. | Train. | Val. | Train. | Val. | Train. |
| $1 \times 10^{-1}$ | 0.693<br>0.001 | ± 0.591<br>0.059 | ± 0.430<br>0.093 | ± 0.711<br>0.070 | ± 0.533<br>0.029 | ± 0.733<br>0.058 |
| $1 \times 10^{-3}$ | 0.579<br>0.080 | ± 0.123<br>0.104 | ± 0.795<br>0.093 | ± 0.983<br>0.029 | ± 0.800<br>0.087 | ± 0.983<br>0.029 |
| $1 \times 10^{-5}$ | 0.697<br>0.003 | ± 0.686<br>0.004 | ± 0.333<br>0.000 | ± 0.333<br>0.000 | ± 0.500<br>0.000 | ± 0.500<br>0.000 |

### Attended spectra instances

To gain insight into the model’s decision-making process, we analysed the attention weights assigned to the spectra within the validation dataset. This attention mechanism is a key feature for interpretability, as it reveals which spectral features and spatial regions the optimised model considered most significant for classification. First, the validation dataset was processed by the fully trained model, and the attention weight for each spectral instance was extracted. For each hyperspectral image, these weights were aggregated and then normalised to a range of 0 to 1. To visualise the spatial distribution of these important regions, the normalised attention weights were reshaped into two-dimensional ’attention maps’ matching the original chemical image dimensions (480 x 480 pixels). These maps were then overlaid onto a greyscale image representing the tissue morphology region on the slide. This morphological base image was generated by calculating the Area Under the Curve (AUC) for each spectrum, which provides the total integrated signal intensity at each pixel (*i,j*). The AUC was calculated using the following integral, where *P(ν)* is the spectral intensity at wavenumber, *ν*:

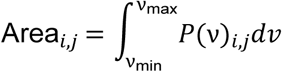

Finally, for more targeted downstream analysis, we identified and isolated the most influential spectra as determined by the model. From each chemical image in the validation set, the spectra corresponding to the top 5% of attention weights were filtered and aggregated. This process created a refined dataset comprising only the spectra that the model deemed most salient, providing a high-value subset for subsequent investigation.

### Interpreting model predictions with SHAP

To gain a deeper insight into the spectral features driving the model’s predictions, we analysed the data using SHapley Additive exPlanations (SHAP). This analysis was performed with the *shap* Python library (version 0.48.0) (22). The primary objective was to identify the key wavenumbers that have the most significant impact on distinguishing between classes, thereby enhancing model interpretability and facilitating chemical inference from its outcomes. For this task, we specifically employed SHAP’s GradientExplainer, an approach optimised for deep learning models. This explainer was applied to the validation dataset to calculate feature importance scores. The resulting SHAP values were then used to generate a summary plot for each class ranking the wavenumbers based on their overall importance to the model and illustrate not only the magnitude, but also the direction of each feature’s effect. This will show whether a high absorbance at a specific wavenumber pushes the prediction towards or away from a particular class. It also allows for the direct identification of the key spectral bands that underpin the model’s performance.

### UMAP clustering

To determine if the top spectra identified by the model’s attention mechanism possess biological relevance, we investigated whether they could be separated based on treatment type. This analysis was performed using Uniform Manifold Approximation and Projection (UMAP), implemented via the *umap-learn* Python library (version 0.5.7) (23). UMAP is a powerful non-linear dimensionality reduction technique used to project high-dimensional data into a low-dimensional embedding, making it easier to visualise data structure and class separability (23). Prior to UMAP analysis, these input spectral features were scaled using a Standard scaler from the *scikit-learn* library [20]. This pre-processing step normalises the distribution of each feature, ensuring that the intensity scaling is consistent across all wavenumbers and improving the robustness of the subsequent distance-based calculations in UMAP.

A systematic parameter search was conducted to find the optimal UMAP configuration that could most effectively cluster the spectra according to their treatment class (FOLFOX, ABT-199, FOLFOX with ABT-199, or vehicle control). The key parameters explored included the number of neighbours (*n_neighbors*, ranging from 5 to 80), the minimum distance between embedded points (*min_dist*, ranging from 0.1 to 0.5), and various distance metrics (Euclidean, Cosine, and Manhattan). For each set of parameters, a two-dimensional UMAP embedding of the spectra was generated. This low-dimensional representation was then clustered using the k-means algorithm, with the number of clusters (*k*) set to seven, corresponding to the number of distinct treatment labels. The performance of each UMAP configuration was quantified by evaluating the agreement between the k-means cluster assignments and the ground-truth treatment labels. This was measured using the Normalised Mutual Information (NMI) score, a metric that assesses the similarity between the clustering and the ground truth class membership (24,25). The NMI between the true labels (*U*) and the k-means cluster assignments (*V*) is calculated as:

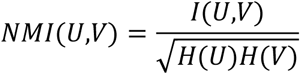

where, *I(U,V)* represents the mutual information between the two assignments, while *H(U)* and *H(V)* are their respective entropies. The optimal UMAP parameters were selected based on the configuration that yielded the highest NMI score, as this indicates the clearest and most meaningful separation of the treatment groups in the low-dimensional space.

### Correlation study

An additional correlation analysis was conducted to investigate the relationship between spectral data and biological endpoints. We aimed to investigate whether the changes of the pro-apoptotic proteins Puma and Bim, key mediators of chemotherapy-induced apoptosis and evaluated in the original study [10], lead to detectable spectral bands, using selected wavenumber and ratios. We extracted four protein-related spectral bands: the integrated intensity of the Amide I (1600 - 1700 cm⁻¹), Amide II (1540 - 1560cm⁻¹), and Amide III (1200 - 1350cm⁻¹) bands, and the Amide I/II peak ratio (∼1658/1544 cm⁻¹). Additionally, a series of spectral band ratios were also calculated to assess the relative content of proteins to other key biomolecules, including lipids and nucleic acids. These included ratios of Amide I to Amide II, DNA, RNA, and lipids, providing further insight into the overall biochemical changes within the cells. Details on the ratio’s used and their wavenumber bands are available at the Supplementary section (refer to **Error! Reference source not found.**). Then, using relative protein expression data acquired in the original PDX study (10), a Pearson correlation analysis was conducted independently for each chemotherapy sensitivity group (CRC0076 and CRC0344) to link the spectral metrics with apoptotic protein levels using correlation function available in Pandas python module (version 2.2.2).

## 3. Results and Discussion

### Model architecture parameter optimisation

Our investigation into the optimal model architecture began by examining the training dynamics of models without residual connections. By analysing the mean of F1-score and loss curves over 20 epochs and three trial runs, we observed starkly different behaviours between the ’shrinking’ (half) and ’expanding’ (double) network configurations (see **Figure 2**). The training curves for the ’half’ expansion strategy demonstrate a stable learning process where for these models, the validation F1-scores (dashed lines) follow the training F1-scores (solid lines), indicating that the models were generalising to the unseen data. The corresponding loss curves of hidden dimension 128 corroborate this positive trend, with both training and validation losses decreasing in tandem, signifying effective learning without significant overfitting except for hidden dimension 64 and 256 where the loss validation losses stay pretty constant throughout the epochs.

**Figure 2.**
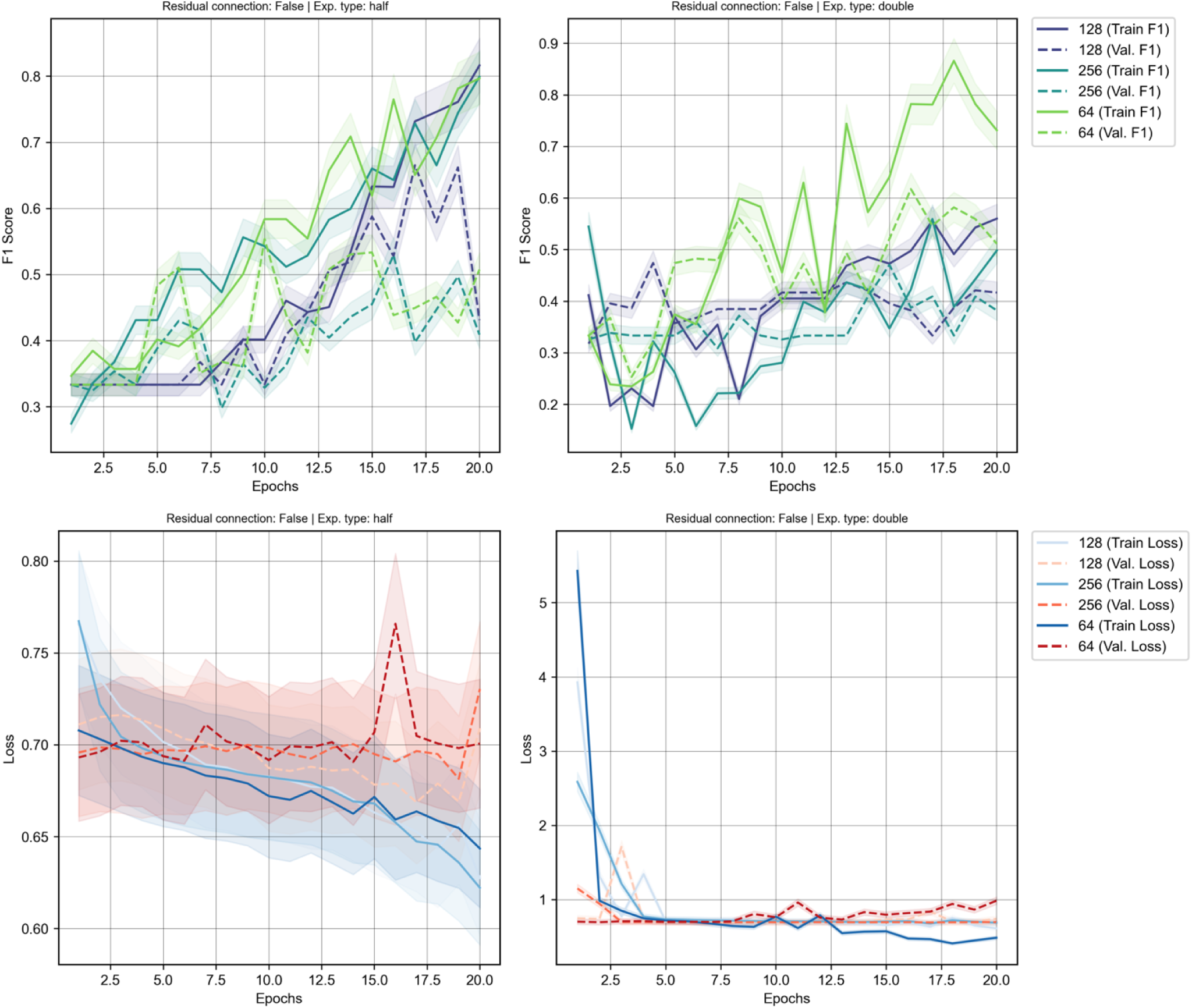
Training results for model architecture optimisation without residual connection using different hidden number of nodes accumulated over three trial runs. Left panel show ‘half’ expansion type while eight panel show ‘double’ expansion type. Top panel is the training and validation F1-score and bottom panel is the loss values accumulated during training phase. Shaded regions are the standard deviation x 0.05 scale factor.

Conversely, the training dynamics for the ’double’ expansion strategy reveal different patterns where the F1 scores slightly worsen at the beginning of the training epochs before slowly improving over time. Different parameters also behave differently in this expansion strategy where diverging patterns can be observed to either significantly improve the overall F1 scores (hidden dimension 64) or cause them to remain stagnant (hidden dimension 128 and 256) when compared to the ’half’ expansion model type in the validation dataset. This divergence is a definitive sign that the rapidly increasing complexity of the expanding network to analyse or focus more on the noise in the data rather than learn generalisable, predictive features.

These training dynamics are reflected in the overall performance landscapes (see **Figure 3**), which visualise an aggregate of the mean validation F1-scores from multiple runs across the hyperparameter grid. The landscape for the ’half’ strategy reveals a broad, stable region of good performance, peaking at an average F1-score of approximately 0.77 for a configuration of 2 blocks and an initial hidden dimension of 256. In contrast, the landscape for the ’double’ strategy is more volatile, with performance dropping off sharply for deeper models, reinforcing the conclusions drawn from the unstable training curves. Interestingly, while the average performance favours the ’half’ strategy, examining the best result from all training run presents a more nuanced picture. The highest F1-score achieved by any ’double’ configuration was 0.829, obtained with a single block and a 128-node hidden dimension. However, the overall best-performing individual model was a ’half’ configuration (using 64 hidden nodes), which reached a remarkable F1-score of 0.950 in one of its trials. This highlights a key trade-off: while certain ’double’ configurations show high potential, their performance is inconsistent. The ’half’ architecture not only produced the single best result but also demonstrated significantly more stable performance on average, making it the more dependable and promising architectural choice for this task.

**Figure 3.**
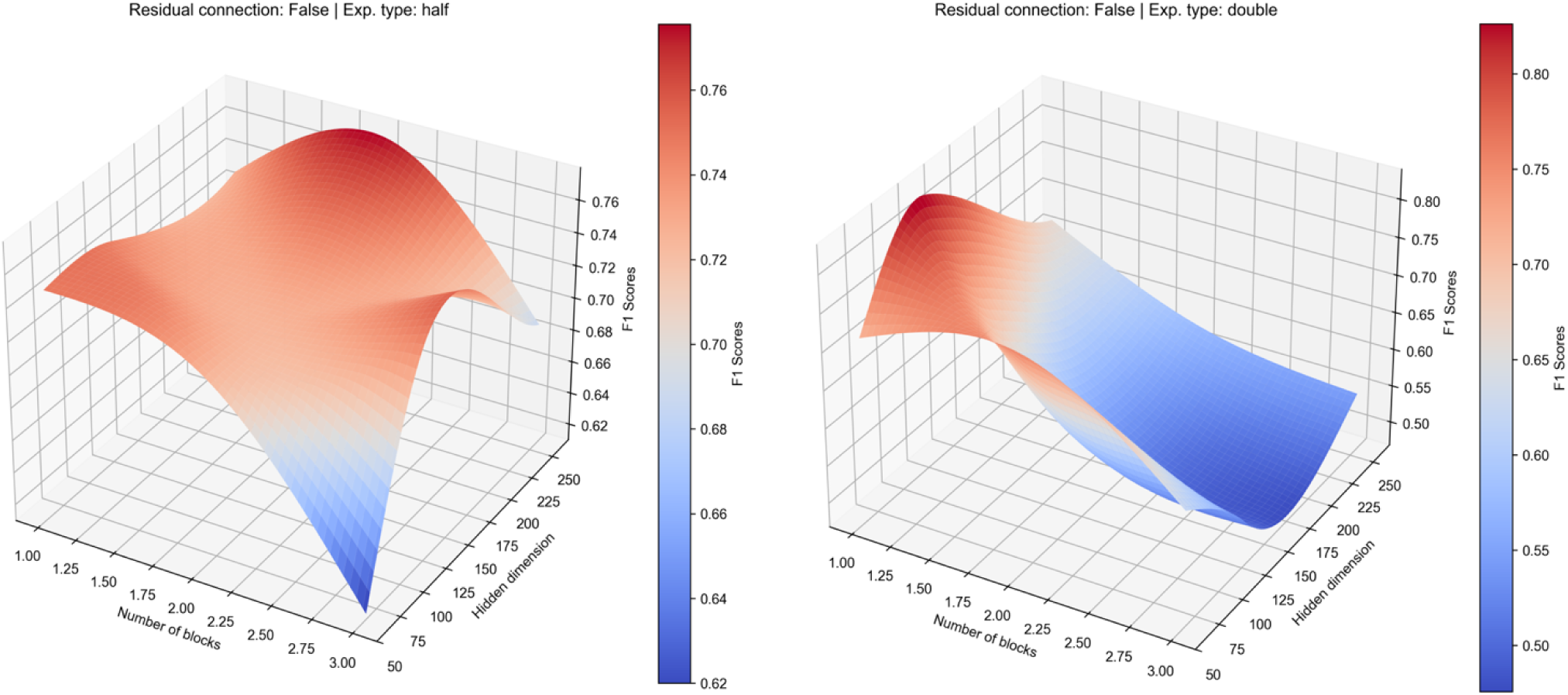
Average performance landscape of the validation dataset showing how different model architecture parameters impact the model prediction without residual connection. Left panel show ‘half’ expansion type and right panel show ‘double’ expansion type.

### Model architecture optimisation with residual connections

Following the initial architectural optimisation, we investigated the impact of incorporating residual connections into the model. These connections were designed to improve gradient flow and mitigate the vanishing gradient problem in deeper networks, theoretically yielding a significant improvement in both model performance and training stability across the board. The ’half’ (shrinking) architecture, which was already the top performer, saw further gains with the addition of residual connections when comparing the train F1-score (see **Figure 4**). The training curves reveal an even more stable and faster learning process, with the validation F1-score tracking the training scores closely, indicating good model generalisation. However in terms of the validation F1-score, the performance of this configuration slightly under performs in comparison with the non-residual model architecture. A deeper analysis of the loss values during training suggested that the poor validation F1-scores were due to the overfitting on the training set, as the loss values on the validation dataset quickly diverge at the very early of the training epochs. This suggests that for an already stable architecture, the addition of residual connections does not provide a definitive advantage in generalisation for this specific task.

**Figure 4.**
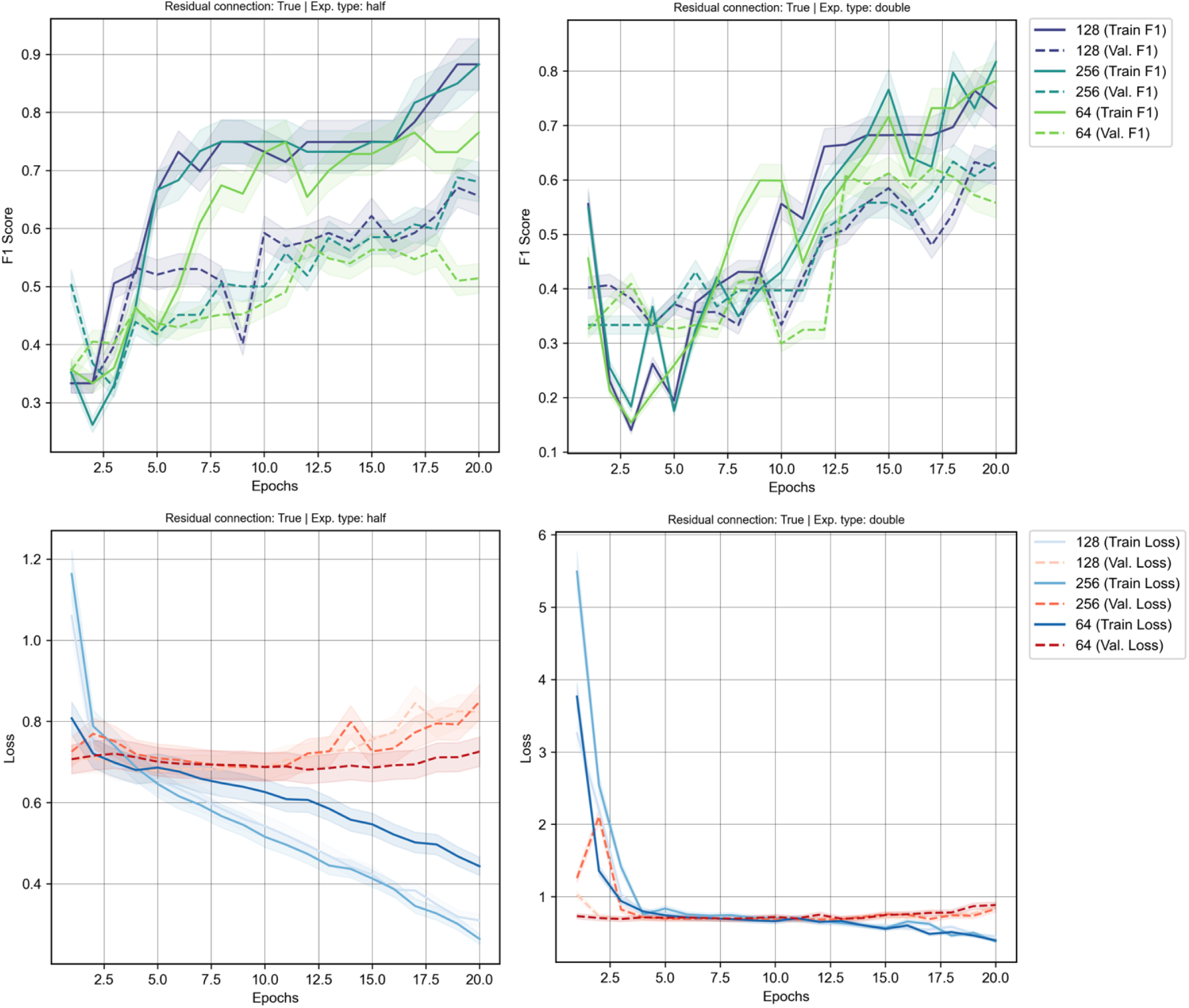
Training results for model architecture optimisation with residual connection using different hidden number of nodes accumulated over three trial runs. Left panel show ‘half’ expansion type while right panel show ‘double’ expansion type. Top panel is the training and validation F1-score and bottom panel is the loss values accumulated during training phase. Shaded regions are the standard deviation x 0.05 scale factor.

The most dramatic effect was observed in the ’expanding’ (double) architecture, which was stabilised by the inclusion of these connections (see **Figure 4**). The model’s generalisation improved significantly, converging faster and achieving a better validation F1-score compared to the same configuration without residual connections. This indicates that when a model has a very high number of parameters, residual connections are crucial for stabilising the learning process. The loss values, while starting at a higher level, became more stable over time and the gap between training and validation F1-scores was significantly reduced. This demonstrates that residual connections enable more effective gradient flow through the network, preventing the model from exclusively memorising the training data and instead forcing it to learn more robust, generalisable features. While its peak average performance, with an F1-score of around 0.72, still fell short of the ’half’ architecture, the model was no longer failing to generalise. This stabilising effect is visually confirmed in the performance landscape, which transformed from a volatile surface to a much smoother and more predictable configuration (see **Figure 5**).

**Figure 5.**
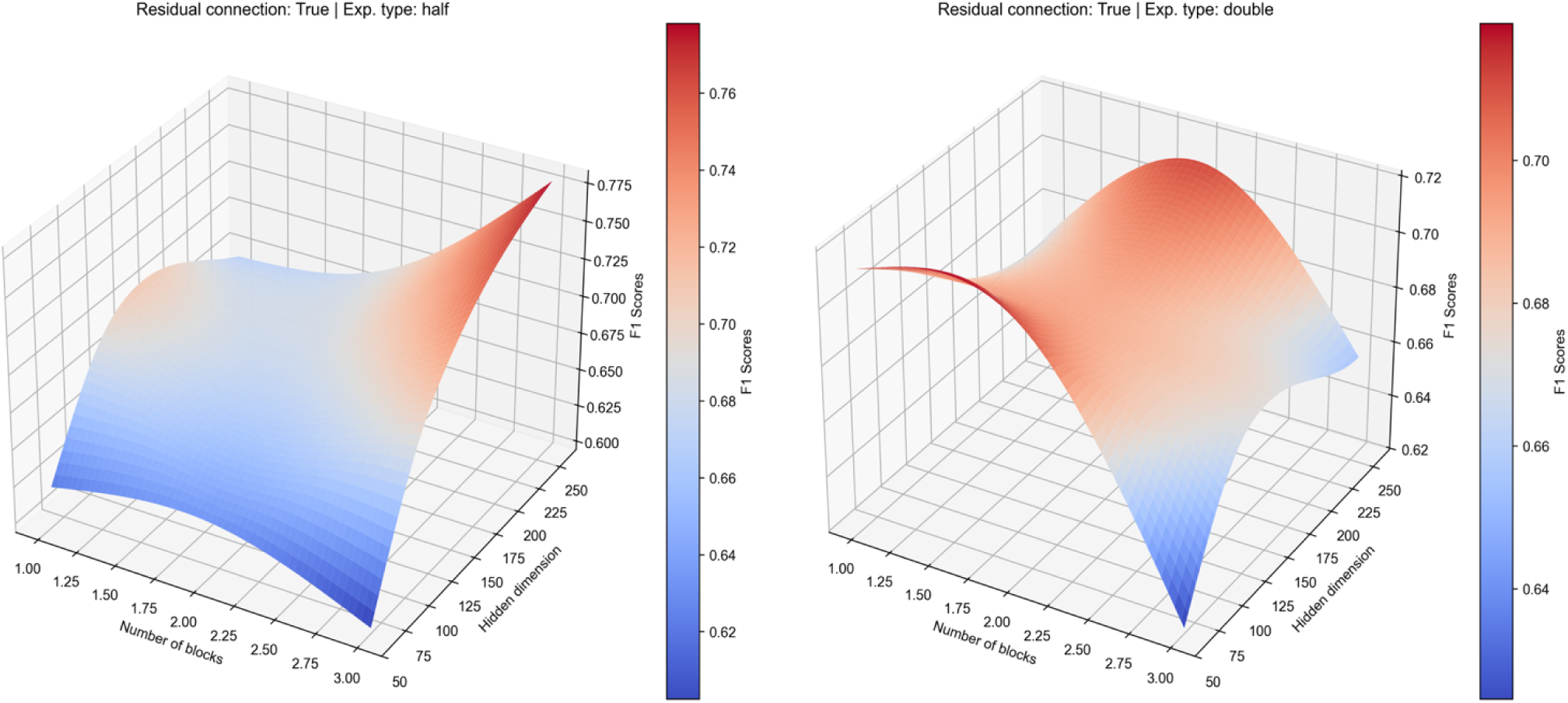
Average performance landscape of the validation dataset showing how different model architecture parameters impact the model prediction with residual connection. Left panel show ‘half’ expansion type and right panel show ‘double’ expansion type.

In conclusion, our comprehensive architectural evaluation revealed the clear superiority of the ’half’ (shrinking) strategy over the ’expanding’ approach. While residual connections proved essential for stabilising the otherwise volatile ’double’ models, they did not consistently improve the performance of the already robust ’shrinking’ models. The selection of the final model was based on identifying the configuration that achieved the highest peak performance across all individual training runs, prioritising demonstrated potential over average performance. Based on this criterion, the optimal model was determined to be a non-residual architecture with 2 blocks, a ’half’ expansion strategy, and an initial hidden dimension of 64 nodes (N_B_=2, Residual=False, Exp. type=’half’, N_HLN_=64). In a single training run, this specific configuration achieved the highest validation F1-score of 0.950 across all tested parameters and executions. This outstanding result suggests that for this dataset, a more compact and simpler architecture was best suited to find an optimal solution without succumbing to overfitting. Therefore, this top-performing model was selected for all subsequent analyses.

### Learning rate parameter optimisation

Once the optimal model architecture was established, the final step in our hyperparameter tuning was to optimise the learning rate for the Adam optimiser. The learning rate is a critical parameter that controls the step size the model takes during gradient descent. Selecting an appropriate learning rate is crucial for ensuring efficient convergence to a good minimum on the loss landscape, thereby preventing unstable training or premature termination in a suboptimal solution. To identify the optimised setting, we evaluated three distinct learning rates spanning several orders of magnitude: a high rate (1x10⁻¹), a moderate rate (1x10⁻³), and a low rate (1x10⁻⁵). Each was tested on the chosen architecture, with the resulting F1-score, accuracy, and loss values recorded to determine the most effective option (see bookmark6 **self-reference.**).

The results clearly demonstrated the sensitivity of the model to this parameter. The high learning rate of 1x10⁻¹ resulted in unstable training, characterised by high loss and poor performance metrics, as the optimiser likely overshot the optimal solution repeatedly. Conversely, a low learning rate of 1x10⁻⁵ led to excessively slow convergence, failing to achieve a competitive F1-score within the training epochs. A moderate learning rate of 1x10⁻³ provided the best balance, achieving the lowest loss and the highest F1-score and accuracy. This indicates that it was effective at navigating the loss landscape, allowing the model to converge efficiently and stably to a high-quality solution. A learning rate of 1x10⁻³ was therefore selected as the optimal value for the Adam optimiser. This rate enabled robust and efficient training of our final model architecture.

With all architectural hyperparameters and learning rate value finalised, the optimal model was retrained for an extended duration to ensure full convergence. The selected configuration; a non-residual, shrinking network with 2 blocks, 64 initial hidden nodes, and a learning rate of 1x10⁻³ was used for training up to 50 epochs. During this final training run, model checkpoints were saved each time a new highest validation F1-score was achieved. This strategy ensures that the selected model represents the point of peak generalisation performance. The best-performing checkpoint yielded a final validation F1-score of 0.9499. This highly performance model was saved and used for all subsequent analysis and interpretation in this study.

### Uncovering salient tissue regions using attended spectral instances

A key advantage of our Multiple Instance Learning (MIL) framework is its inherent interpretability due to the integration of an attention mechanism. This mechanism allows the model to sift through hundreds of thousands of spectral instances within each hyperspectral image and assign an ’attention weight’ to each one, signifying its importance for the final classification. By extracting these weights, we can gain direct insight into the model’s decision-making process, effectively asking the model which specific spectra it found most informative for distinguishing between the CRC0076 and CRC0344 classes. We generated spatial attention maps for each image in the validation dataset. As shown in the representative examples in **Figure 6**, these maps highlight the specific pixels (and their corresponding spectra) that received the highest attention weights. For the CRC0076 class, the model appears to focus on more discrete, localised regions within the tissue, whereas for the CRC0344 class, the highly attended pixels seem to be more diffusely distributed throughout the tissue structure. This visualisation powerfully demonstrates the model’s ability to identify salient regions without any prior spatial annotation or manual tissue segmentation.

**Figure 6.**
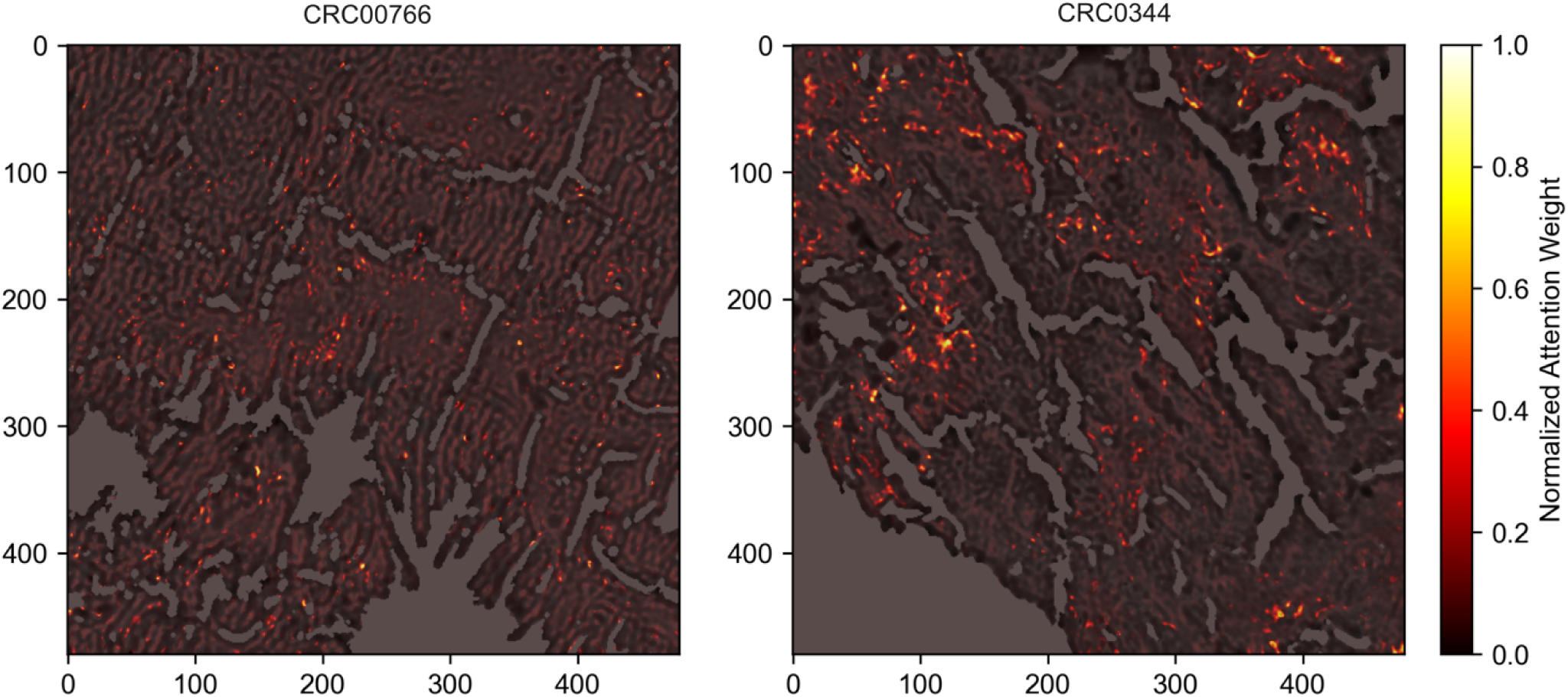
Attention maps from two representative samples from each group (left panel is CRC0076 and right panel CRC0344) obtained from normalised attention weight generated by the MIL mechanism in the model. Grayscale image background is the tissue area under the curve projection while the coloured regions are the intensity of the normalised attention weights.

This weakly supervised approach offers two significant advantages over traditional methods. Firstly, it circumvents the need for laborious and potentially subjective manual annotation of specific tissue regions, a common bottleneck in digital pathology. The model learns to find the relevant areas on its own. Secondly, our MIL approach efficiently handles the immense dimensionality of hyperspectral data. By treating each image as a ’bag’ of 1D spectra rather than a 3D data cube (480x480 pixels x 213 channels), we avoid the massive memory consumption and significantly longer training times associated with 2D or 3D convolution-based models. Ultimately, this attention mechanism provides more than just a visual aid; it acts as a potential data-driven filter for biomarker discovery and could be useful for application to other spatial-biology imaging modalities.

### Model prediction analysis using SHAP

To understand the biochemical basis for the model’s successful classifications, we employed SHAP’s Gradient explainer. This technique allows us to peer inside the “black box” of the complex deep learning model using calculated gradients and identify which specific spectral features were most influential in its decision-making process. The power of this analysis is demonstrated by our MIL strategy, which required the model to autonomously identify these key features from a vast pool of over 200,000 individual spectral instances within each hyperspectral image. The SHAP analysis revealed that the model’s predictions are driven by distinct, biochemically relevant spectral regions (see **Figure 7**). The two most influential areas for classifying both CRC0076 and CRC0344 were the phosphate stretching region (ν-PO₄³⁻) around 965 cm⁻¹ and the carbonyl stretching region (ν-C=O) between 1710-1745 cm⁻¹, associated with lipids and RNA. The high SHAP values across both classes in these regions indicate that the model’s fundamental logic is built on detecting core differences in the state of nucleic acids and lipids. It is known that chemotherapy alters the lipid profile of cancer patients, affecting lipoproteins such as very-low-density lipoprotein (VLDL) and low-density lipoprotein (LDL), as well as apolipoproteins including apolipoprotein A1 (26,27). In retrospect, we also compared this result with the findings from the patient sample source. mCRC frequently has several somatic mutations that reduce response to therapy (16). This comprises *KRAS* mutations, *TP53* mutations, *APC* mutations, *BRAF* mutations, and *NRAS* mutations. CRC0076 harbours *APC-R1114*, *TP53-R273H* and *KRAS-G12V* mutations, while CRC0344 shows *APC-E1309fs*4*, *TP53-R248W* and KRAS-G12D mutations. While there was a comparable mutation spectrum for these four genes, small differences in their biological effects or the presence of other somatic mutations or epigenetic changes may account for the nucleic acid spectral band contributions to the model classification.

**Figure 7.**
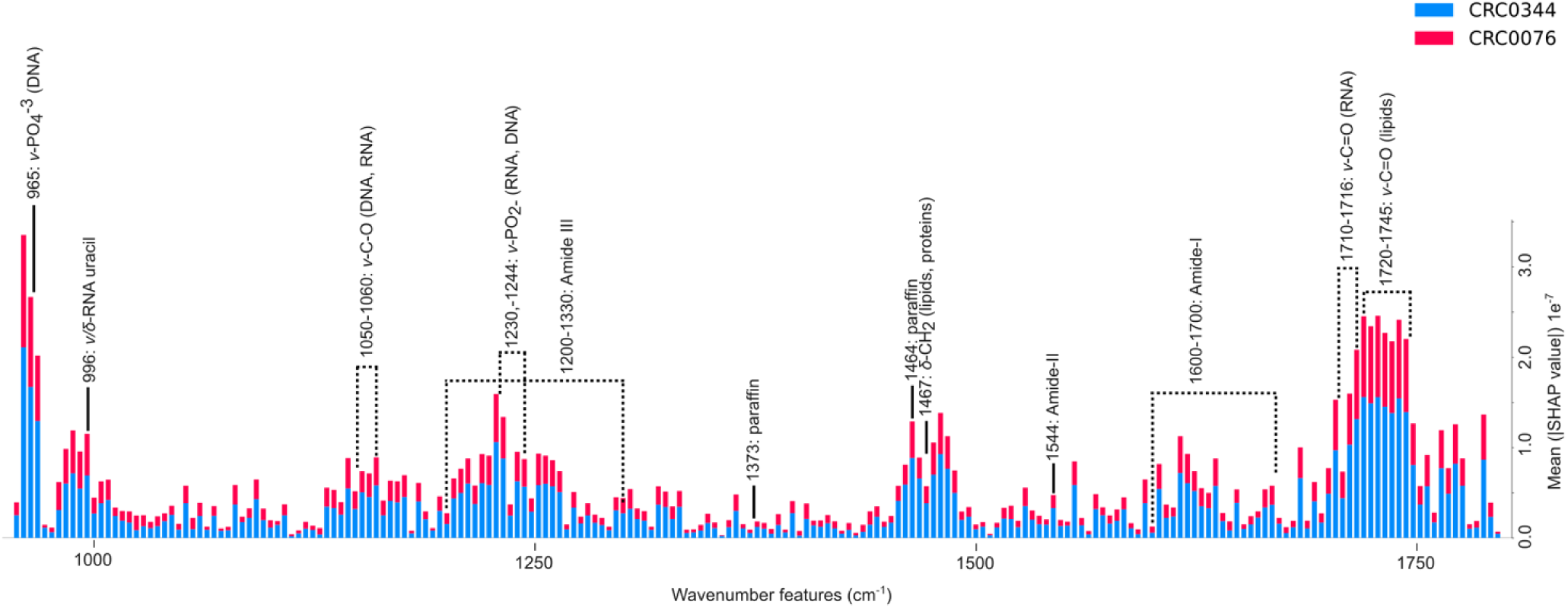
The important wavenumber features based on the mean SHAP values of the two different classes that contribute to the model classification decision. The black line indicates specific location of the molecular vibrations within the IR range while the dotted line indicates its range. ν (nu) represent stretching vibration of the atomic bonds (bond length) while δ (delta) represent bending vibration (bond angle i.e; scissoring, rocking, wagging and twisting).

Beyond these dominant regions, the model also moderately utilised spectral information from protein structures. The broad Amide I and Amide III bands, both sensitive to protein secondary structure, consistently showed high feature importance. Furthermore, the model demonstrated high specificity by assigning significant importance to the peak at 996 cm⁻¹, attributed to stretching and bending vibrations of the RNA uracil base (ν/δ-RNA uracil), highlighting its ability to discern signals beyond the general phosphate backbone. This suggests that the fundamental differences between the two tissue groups are primarily associated with nucleic acids, followed by lipids and then proteins from spectral information of the chemical images generated by IR spectroscopy. Conversely, the contribution of the Amide I region may indicate changes in protein folding or aggregation states, which is a prevalent mechanism by which chemotherapy induces cellular cytotoxic stress to destroy cancer cells (reviewed in (28)).

A key and novel finding of this study is how the model handled sample and experimental artifacts. The peak at 1464 cm⁻¹ and 1373 cm^-1^ correspond to the characteristic of a paraffin wax, was assigned a relatively moderate to very low importance value by the model. Despite the potential for high absorbance from this contaminant in the raw spectra (see **Figure 8**b), the model was intelligent enough to learn that this feature was irrelevant for distinguishing the biological classes. This demonstrates a sophisticated ability to differentiate endogenous biochemical signals from exogenous contaminants, a significant advantage of this data-driven approach. These results powerfully demonstrate the advantage of using the whole spectrum as input for a deep learning model. Rather than relying on engineered features like principal components (29), which can obscure subtle yet important peaks, our MIL approach allows the model to discover novel and specific spectral biomarkers directly from the data, while simultaneously learning to ignore irrelevant information.

**Figure 8.**
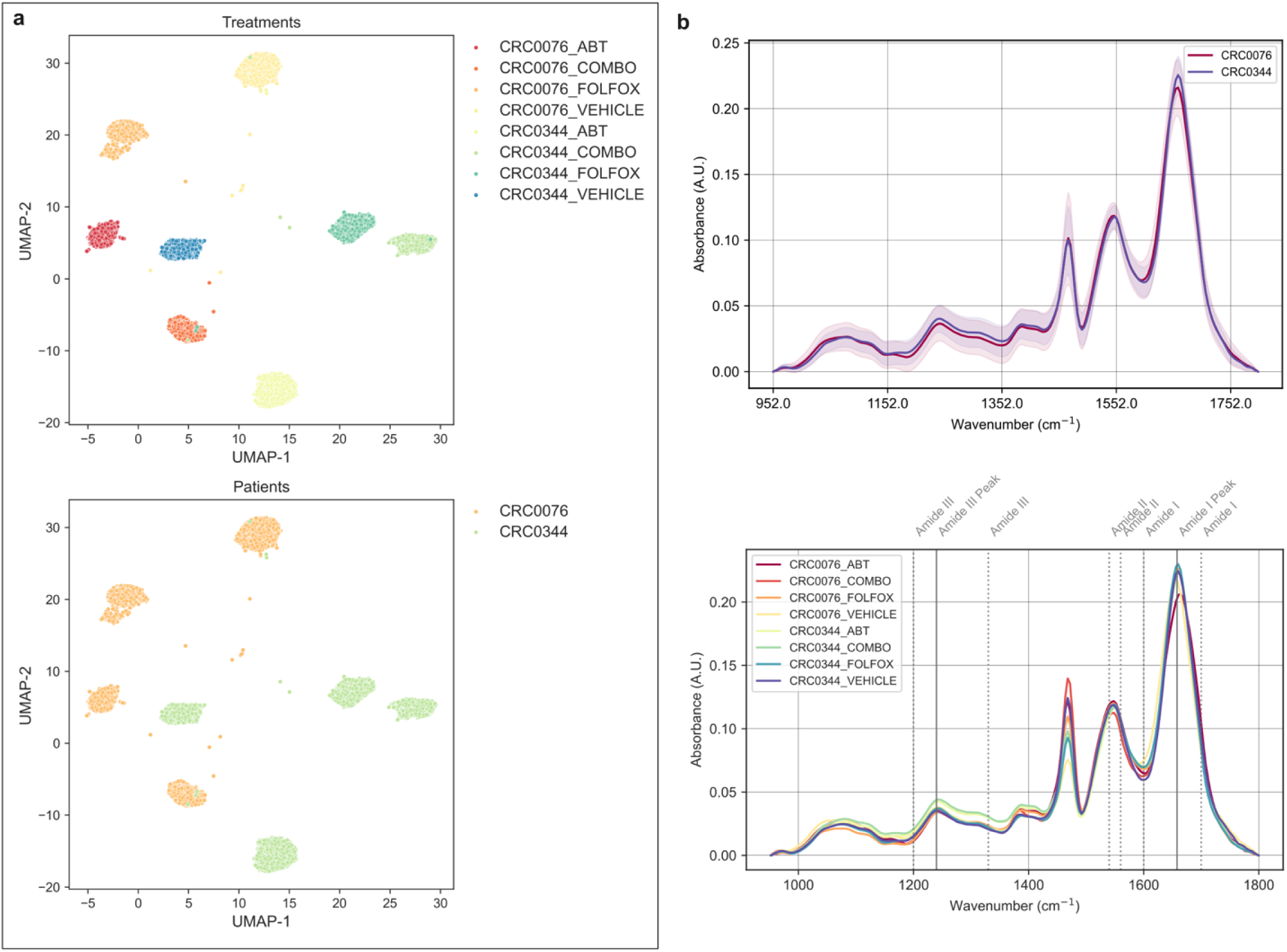
Biological relevance of the most highly attended spectral instances. a) UMAP clustering of the top-attended spectra, illustrating distinct groupings according to treatment type (top panel) and classification label (bottom panel). b) Mean absorbance profiles of the top-attended spectra, shown by classification group (top panel) and treatment type (bottom panel), with annotated protein-related wavenumber bands. Solid grey lines mark specific band positions, while dotted lines indicate the boundaries of the corresponding spectral regions.

### Clustering of salient spectra by treatment pattern

After confirming that our model could identify salient spectra for classifying patient-derived xenograft (PDX) models, we sought to determine if these same spectra contained fine-grained biochemical information related to specific chemotherapy treatments. In the previous study, each PDX model (CRC0076 and CRC0344) had been subjected to different treatment strategies; FOLFOX, the BCL-2 antagonist ABT-199 (ABT199), a combination of both (COMBO), or a vehicle control. We hypothesised that the spectral instances selected by the attention mechanism as important for patient classification might also harbour unique signatures corresponding to these treatments. To investigate this, we applied the non-linear dimensionality reduction technique, Uniform Manifold Approximation and Projection (UMAP), to the collection of top-attended spectra. Following a systematic parameter optimisation, we found that a configuration using a ’euclidean’ distance metric, a number of neighbours of 80, and a minimum distance of 0.25 yielded the best separation of treatment groups. This optimal configuration achieved a near-perfect Normalised Mutual Information (NMI) score of 0.994, confirming an exceptionally strong correlation between the UMAP clusters and the actual treatment labels.

The resulting UMAP embedding, shown in **Figure 8**a (top panel), visually confirms this outstanding separation. Each treatment group forms a distinct and tight cluster, which is itself nested within the patient class. This powerful result demonstrates that the spectral instances prioritised by our model are not only sufficient for robust patient-level classification but are also rich with detailed information capturing the specific molecular changes induced by each therapy. The model, despite being trained only to distinguish CRC0076 from CRC0344, has implicitly learned to focus on spectra that are also sensitive to treatment effects.

To explore the biochemical basis for this separation, we calculated and plotted the mean spectrum for each distinct treatment group (**Figure 8**b, lower panel). The most prominent differences between the spectra were observed in the protein-associated regions, particularly the Amide III (1200-1300 cm⁻¹), Amide II (∼1550 cm⁻¹), and Amide I (∼1650 cm⁻¹) bands. The spectra from the CRC0344 model treated with ABT-199 (both alone and in combination with FOLFOX) display a distinct increase in absorbance in the Amide I and Amide III regions in comparison to the other treatment groups. This spectral signature is highly significant for this context, as alterations in amide bands often reflect changes in protein secondary structure (30) and are frequently associated with cellular apoptosis (programmed cell death). Given that ABT-199 is an apoptotic-inducing agent, this finding provides compelling biochemical evidence that our model successfully identified the spectral footprint of the drug’s mechanism of action directly within the tissue. As reported by O’Farrell et al (10), this CRC0344 group were also more susceptible to the chosen treatment regimen in comparison to the CRC0076.

Taken together, these findings serve as a crucial validation of the features learned by the attention mechanism. The clear separation of treatment groups by unsupervised clustering, and more specifically, the direct correlation between the ABT-199 spectral signature and known pro-apoptotic protein changes, confirms that the attended spectra are not arbitrary. Instead, they represent biochemically significant events within the tissue, demonstrating that our model successfully distilled complex cellular responses into a set of highly informative spectral features.

### Correlation of salient spectra with pro-apoptotic protein expression markers

To validate the biological relevance of the spectral features identified by our model, we sought to connect them to the experimental data from the initial study. This correlation analysis section aimed to correlate key protein spectral markers derived from IR spectroscopy (Amide I, Amide II, and the Ratio Amide I/II) and other ratios of protein to cellular components (i.e. Amide I/Lipid, Amide I/DNA, Amide I/RNA and others) with the expression levels of the pro-apoptotic proteins Bim and Puma, which were quantified previously via Western blot (10). This approach allows for the direct investigation of treatment-induced biochemical changes and a comparison of the chemical information changes in both CRC0076 and CRC0344 PDX model.

Analysis of the marker distributions revealed fundamentally different apoptotic states between the two models (**Figure 9**a). A key strength here is the integration of two distinct experimental modalities; we are comparing targeted relative protein expression from Western blotting with attended spectral absorbance from IR spectroscopy. Finding a correlation between these disparate measurements is significant, as it suggests this rapid, label-free spectral technique directly reflects specific molecular events. Indeed, the CRC0076 model exhibited high baseline expression of both Bim and Puma in the vehicle-treated group, a state consistent with being “primed for apoptosis”, where pro-apoptotic proteins are held in check by high levels of anti-apoptotic BCL-2. Upon treatment, the levels of these proteins decreased (except for Bim in combination group). O’Farrell et al. proposed that ABT-199 treatment in this high-BCL-2 model causes the release and subsequent degradation of sequestered Puma, confirming target engagement (10). Interestingly, this pattern is also observed in the majority of the protein spectra absorbance and spectral ratios values including protein-to-nucleic acid (e.g., ratio of Amide I/DNA (PO_4_^-^), ratio of Amide I/Nucleic Acids (PO_2_^−^) and protein-to-lipid (e.g., ratio of Amide I/Lipid (C=O)) demonstrating a probably spectral relation with the experimental data (IR peak assignments for cellular spectra were based from (31)). The Amide I to RNA (Uracil) ratio, on the other hand, did not follow the overall trend and instead increased following the treatment. In contrast, the CRC0344 model generally showed low baseline expression of these proteins and spectral absorbance values (except for Amide II band, the ratio of Amide I/ Nucleic Acids (PO_2_^−^) and the ratio of Amide I/Lipid (CO-O-C)), with a notable increase in signal following the treatments.

**Figure 9.**
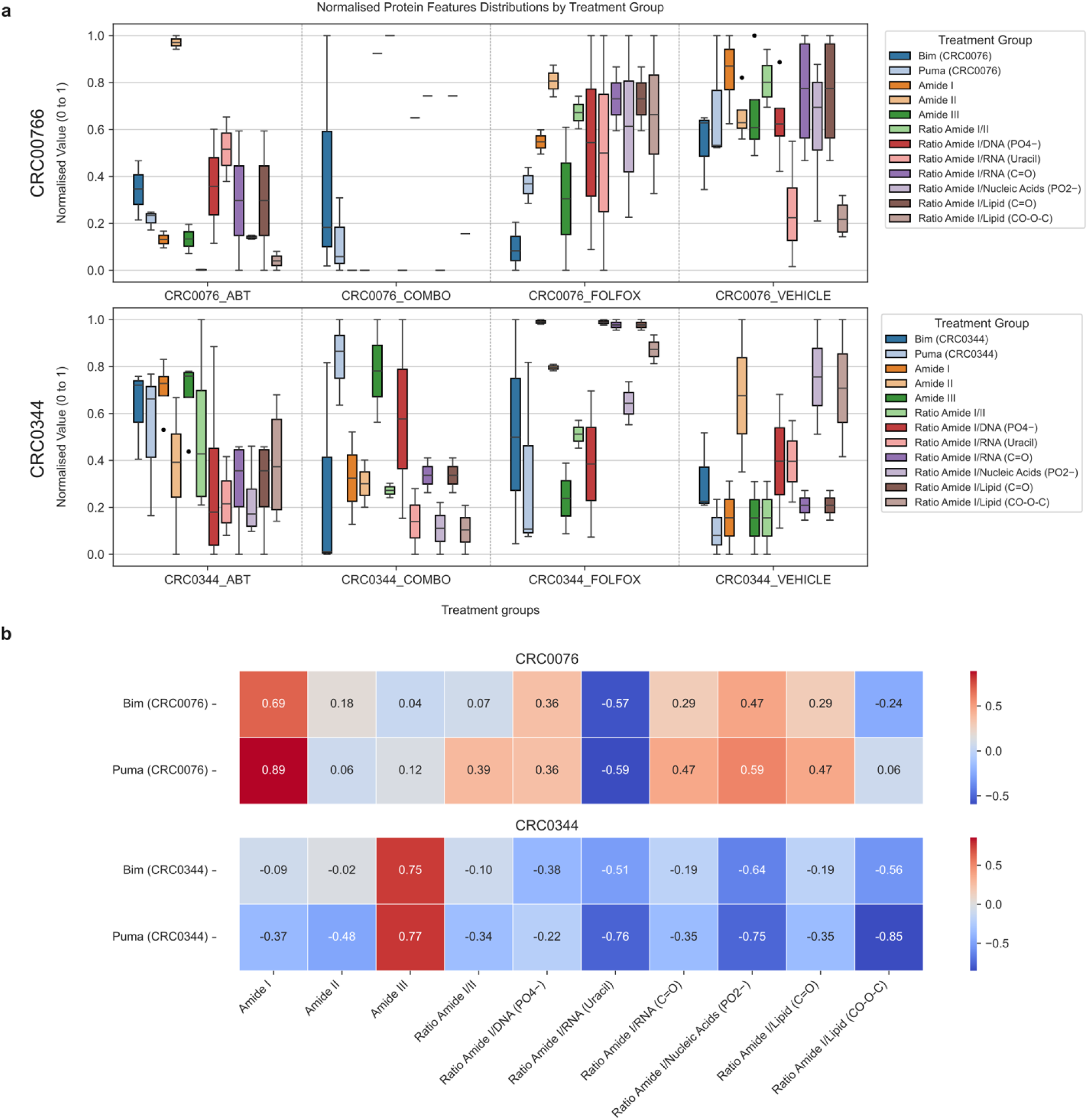
Correlation between selected top-attended spectral features and spectral ratios to the protein expression levels. a) Normalised absorbance values and ratios at specific wavenumbers and corresponding relative protein expression levels for CRC0076 (top panel) and CRC0344 (bottom panel). b) Correlation analysis using Pearson’s correlation between spectral absorbance values and ratios to the relative protein expression levels for CRC0076 (top panel) and CRC0344 (bottom panel).

To validate our initial observation and to quantitatively uncover the relationships between these variables, a correlation analysis was performed (**Figure 9**b) between the spectral bands and the protein expression data. In the CRC0076 model, the Amide I spectral band showed exceptionally strong positive correlations with the expression of both Bim (*r* = 0.69) and the highest is Puma (*r* = 0.89). This link indicates that in this biological context, Amide I intensity is a robust, non-invasive proxy for the status of the Puma- and Bim-mediated apoptotic pathways. The ratio of Amide I/II also supports this, showing a moderate positive correlation with Puma (*r* = 0.39) but not for Bim protein (*r* = 0.07). In contrast, this apoptotic signature was completely different in the CRC0344 model. The strong positive correlations with Amide I were lost, and in some cases, inverted. The Amide III band emerged as the key spectral reporter, showing strong positive correlations with both Bim (*r* = 0.75) and Puma (*r* = 0.77). Furthermore, Puma expression was now negatively correlated with Amide I (*r* = -0.37) and Amide II (*r* = -0.48). This shift underscores a fundamental divergence in the molecular response to therapy between the two patient-derived models. These results validate the biological relevance of the Amide bands as reporters of apoptosis but also demonstrate that their interpretation is highly context-dependent.

Expanding the analysis to the spectral ratios which reflect the balance between macromolecules, reveals an even more complex set of relationships. In the CRC0076 model, the correlations are mixed. A strong positive correlation was found between Bim and Puma to the Ratio Amide I/RNA (C−O) (*r* = 0.57 and 0.59 respectively), reinforcing the link between this apoptotic protein and the overall protein-to-RNA balance. However, this is contrasted by moderate positive and negative correlations between both Bim and Puma and ratios involving lipids and DNA (PO_4_ ^-^). The ratio of Amide I/ DNA (PO_2_ ^-^) is moderately positively associated with both Bim/PUMA suggesting a link to the integrity of the DNA backbone (conformation) or its hydration (32). The observations in the CRC0344 model contrast with these findings, although they are consistent. Here, a uniform negative correlation is observed between the pro-apoptotic protein expression and nearly all calculated ratios. The relationships are particularly strong between Puma and the ratios for Amide I/Nucleic Acids (PO_2_ ^-^) (*r* = -0.75), Amide I/RNA (Uracil) (*r* = -0.76) and Amide I/Lipid (CO-O-C) (*r* = -0.85). This complete inversion of the correlation pattern compared to CRC0076 underscores that the macromolecular shifts during therapy-induced apoptosis are fundamentally distinct in this “non-primed” model. This divergence highlights that while individual amide bands are informative, these spectral ratios provide a deeper, more systemic view of the cellular state, acting as highly sensitive, context-dependent biosignatures of therapeutic response. The identification of distinct biomarker pairs (e.g., Amide I ≡ Puma in CRC0076 vs. Amide III ≡ Bim/Puma in CRC0344) and the ratio of the intensity of Amide I to Lipid or nucleic acids highlights this methodology’s potential for dissecting patient-specific signalling pathways and therapeutic vulnerabilities to chemotherapy. Furthermore, it could deliver new stratification tools for BCL2 antagonists.

## 4. Conclusion

This study successfully demonstrates the development and application of an integrated deep learning-multiple instance learning (DL-MIL) framework for downstream classification tasks with high-content imaging data. In the present instance we apply this approach to the classification of chemotherapy sensitivity using QCL-IR hyperspectral imaging of colorectal cancer PDX models. Through a systematic optimisation process, we identified a robust, non-residual linear neural network architecture capable of distinguishing between chemo-sensitive (CRC0344) and chemo-resistant (CRC0076) models with high fidelity (F1-score=0.95). The MIL attention mechanism proved highly effective, autonomously identifying salient tissue regions from over 200,000 spectral instances per image, thereby circumventing the need for laborious manual annotation. Explainable AI, through the use of SHAP, provided unprecedented insight into the model’s decision-making process as it reveals that the classification was not based on arbitrary patterns but on distinct and biochemically relevant spectral features, primarily related to nucleic acid phosphate backbones, lipid carbonyl groups, and protein amide bands. Critically, this analysis also demonstrated the model’s sophistication in learning to ignore confounding signals from experimental artifacts such as paraffin wax.

Furthermore, we validated the biological relevance of the model-attended spectra through two distinct approaches. Unsupervised UMAP clustering of these spectra successfully stratified the samples not only by their patient-of-origin but also by their specific treatment group, confirming that the learned features were rich with pharmacodynamic information. Secondly, correlation studies established a direct, albeit highly context-dependent, link between these spectral markers and the expression of key apoptotic proteins, Bim and Puma. The discovery of distinct biomarker pairs for each PDX model (e.g., Amide I/Puma for CRC0076 vs. Amide III/Puma for CRC0344) underscores the importance of discovering patient-specific molecular signatures.

In summary, this research presents a powerful, end-to-end pipeline for the label-free, data-driven discovery of spatial biomarkers from complex high-content imaging data by leveraging the entire spatial signature without prior feature engineering. It also highlights the potential of explainable AI-driven spectral histopathology to advance pre-clinical drug efficacy studies and paves the way for developing more precise, patient-specific diagnostic and prognostic spectral-driven tools for biomedical and pre-clinical applications in the future.

## Associated content

Supplementary information: Tables containing list of samples ID used for dataset and spectral ratios used for the correlation study.

## Author Contributions

ADM conceived the study with JHMP and acquired funding. LS and ATB supplied the tissue for analysis, with ATB, ACO and MAJ performing the original PDX study. ATB, JHMP and ACO supplied ground truth experimental data for correlation to chemical image findings. MRR conducted all imaging and the majority of data modelling, with some contributions to coding from RS and AD under the direction of ADM. MRR and ADM wrote and edited the manuscript. All authors contributed to manuscript review.

## Data and Software Availability

All of the spectral data acquired as part of the current study is publicly available on Zenodo at https://doi.org/10.5281/zenodo.17099733. The Linear-MIL model is also available on Github and Zenodo and is at https://doi.org/10.5281/zenodo.17179431.

## Funding Sources / Acknowledgements

ADM, MRR and JHMP were supported by the North-South Research Programme administered by the Higher Education Authority on behalf of the Department of Further and Higher Education, Research, Innovation and Science and the Shared Island Fund (AICRIstart: A Foundation Stone for the All-Island Cancer Research Institute (AICRI): Building Critical Mass in Precision Cancer Medicine https://www.aicri.org/aicristart).

ADM, MRR and JHMP were also financially supported by Research Ireland through its Infrastructure Programme (Grant Agreement No. 21/RI/9787; National Spatial Tissue Profiling (NaSPro) Platform for Precision Medicine: Integration of Chemical, Brightfield and Fluorescence Imaging of Tissues Combined with Single Cell Molecular Profiling). ADM and MRR were also supported by the ADAPT Research Ireland research Centre at TU Dublin. The ADAPT Centre for Digital Media Technology is funded by Research Ireland through its Research Centers Programme and is co-funded under the European Regional Development Fund (ERDF) through Grant # 13/RC/2106_P2.

ATB acknowledges funding from Science Foundation Ireland (Awards 13/CDA/2183 and 20/FFP-P/8884) and the EU Horizon 2020 Health Research and Innovation award ‘Colossus’ (Grant Agreement number 754923).

